# Calcium-vesicles perform active diffusion in the sea urchin embryo during larval biomineralization

**DOI:** 10.1101/2020.08.14.244053

**Authors:** Mark R. Winter, Miri Morgulis, Tsvia Gildor, Andrew R. Cohen, Smadar Ben-Tabou de-Leon

## Abstract

Biomineralization is the process by which organisms use minerals to harden their tissues and provide them with physical support. Biomineralizing cells concentrate the mineral in vesicles that they secret into a dedicated compartment where crystallization occurs. The dynamics of vesicle motion and the molecular mechanisms that control it, are not well understood. Sea urchin larval skeletogenesis provides an excellent platform for investigating the kinetics of mineral-bearing vesicles. Here we used lattice light-sheet microscopy to study the three-dimensional (3D) dynamics of calcium-bearing vesicles in the cells of normal sea urchin embryos and of embryos where skeletogenesis is blocked through the inhibition of Vascular Endothelial Growth Factor Receptor (VEGFR). We developed computational tools for displaying 3D-volumetric movies and for automatically quantifying vesicle dynamics. Our findings imply that calcium vesicles perform an active diffusion motion in both, calcifying (skeletogenic) and non-calcifying (ectodermal) cells of the embryo. The diffusion coefficient and vesicle speed are larger in the mesenchymal skeletogenic cells compared to the epithelial ectodermal cells. These differences are possibly due to the distinct mechanical properties of the two tissues, demonstrated by the enhanced f-actin accumulation and myosinII activity in the ectodermal cells compared to the skeletogenic cells. Vesicle motion is not directed toward the biomineralization compartment, but the vesicles slow down when they approach it, and probably bind for mineral deposition. VEGFR inhibition leads to an increase of vesicle volume but hardly changes vesicle kinetics and doesn’t affect f-actin accumulation and myosinII activity. Thus, calcium vesicles perform an active diffusion motion in the cell of the sea urchin embryo, with diffusion length and speed that inversely correlate with the strength of the actomyosin network. Overall, our studies provide an unprecedented view of calcium vesicle 3D-dynamics and point toward cytoskeleton remodeling as an important effector of the motion of mineral-bearing vesicles.

**Authors summary:** Biomineralization is a widespread, fundamental process by which organisms use minerals to harden their tissues. Mineral-bearing vesicles were observed in biomineralizing cells and play an essential role in biomineralization, yet little is known about their three-dimensional (3D) dynamics. Here we quantify 3D-vesicle-dynamics during calcite skeleton formation in sea urchin larvae, using lattice-light-sheet microscopy. We discover that calcium vesicles perform a diffusive motion in both calcifying and non-calcifying cells of the embryo. The diffusion coefficient and vesicle speed are higher in the mesenchymal skeletogenic cells compared to the epithelial ectodermal cells. This difference is possibly due to the higher rigidity of the ectodermal cells as demonstrated by the enhanced signal of f-actin and myosinII activity in these cells compared to the skeletogenic cells. The motion of the vesicles in the skeletogenic cells, is not directed toward the biomineralization compartment but the vesicles slow down near it, possibly to deposit their content. Blocking skeletogenesis through the inhibition of Vascular Endothelial Growth Factor Receptor (VEGFR), increases vesicle volume but doesn’t change the diffusion mode and the cytoskeleton markers in the cells. Our studies reveal the active diffusive motion of mineral bearing vesicles that is apparently defined by the mechanical properties of the cells.

**Calcium-vesicle diffusion in biomineralization:** 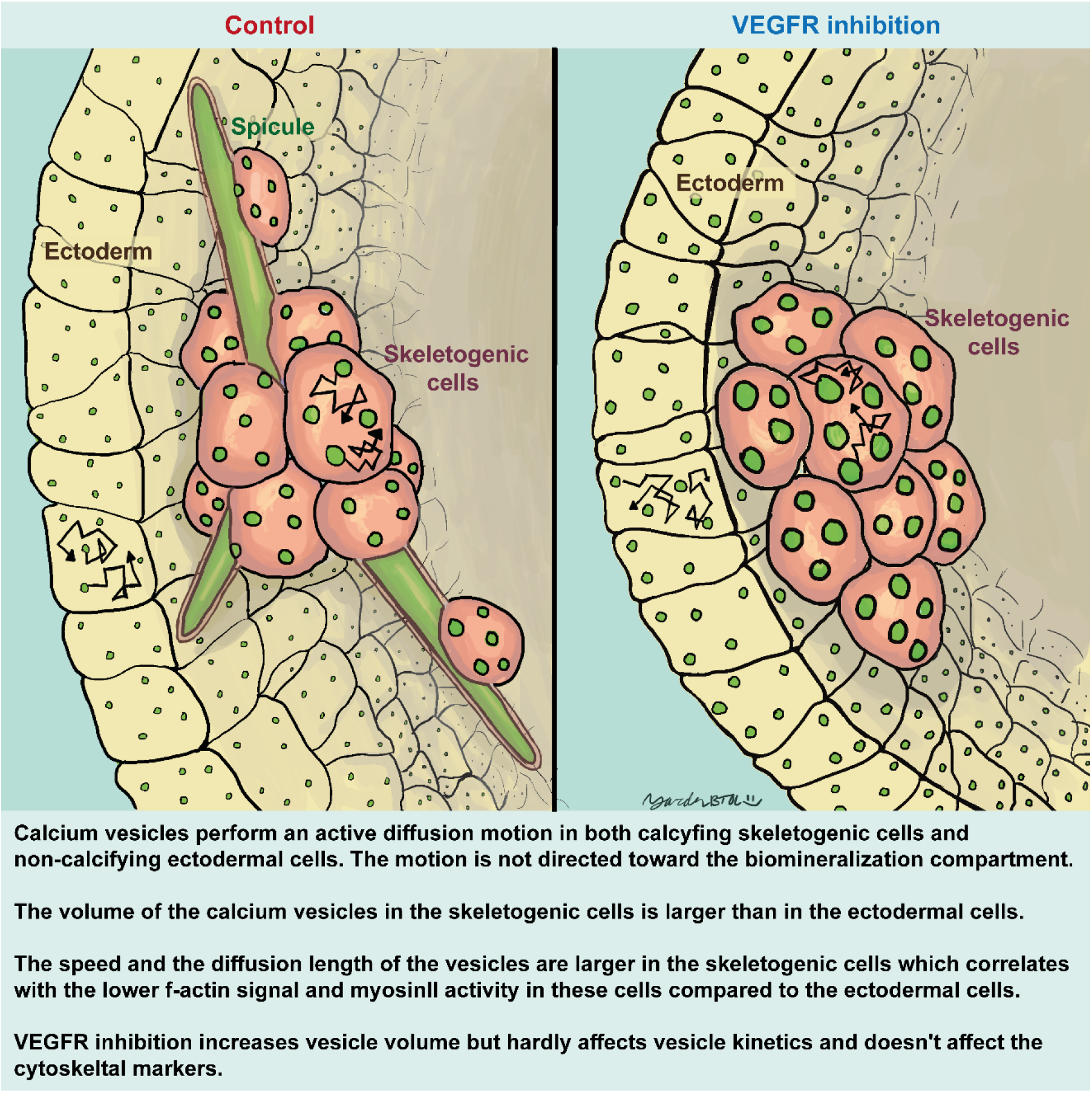

## Introduction

Organisms from the five kingdoms of life use different minerals to harden their tissues and gain protection and physical support [1]. The ability of the cells to control the time and place of crystal nucleation as well as crystal orientation, elasticity and stiffness is beyond the state-of-the art of human technologies [2–4] and inspired the design of biomimetic systems [5–7]. The biomineralized skeletons, teeth and shells, constitute the fossil record that carry the information on the evolution of life on earth. Thus, revealing the cellular and molecular control of the biomineralization process has been the desired goal of both fundamental and applied researchers in the fields of biology, chemistry, geology and material sciences [1, 8–10].

Within the variety of minerals used and phylum specific proteins, a common design of mineral uptake and deposition emerges from various studies: the mineral enters the cells through endocytosis of extracellular fluid [11, 12]. The mineral is then concentrated and maintained in an amorphous phase in intracellular vesicles until the vesicles are deposited into a dedicated compartment where crystallization occurs [13, 14]. The biomineralization compartment provides a highly regulated environment for crystal nucleation and growth. The molecular mechanisms that regulate the trafficking of mineral vesicles and those that control vesicle secretion into the biomineralization compartment, are not well understood.

Sea urchin larval skeletogenesis is an excellent system for investigating mineral uptake, trafficking and deposition within a relatively small (~150μm at gastrula stage) and transparent embryo, which is easy to manipulate [11, 15–22]. The sea urchin skeleton is made of two rods of calcite termed “spicules” generated by the skeletogenic mesodermal cells [19, 23]. The skeletogenic cells fuse and arrange in a ring with two lateral symmetrical cell clusters. In those lateral clusters the skeletogenic cells form a tubular compartment into which calcium-carbonate is deposited to make the calcite spicules [17, 21, 22]. The uptake of calcium from the blastocoel fluid occurs through endocytosis, as was shown through a series of experiments [11, 15, 16]: Electron microscope images of the skeletogenic cells show cell membrane invagination that forms an inner pocket of about 1μm filled with blastocoel fluid [11]. This pocket then separates from the cell membrane and forms an intracellular vesicle with a similar size [11, 15, 16]. Calcium and carbonate are eventually concentrated in the vesicles in the form of amorphous calcium-carbonate (ACC) [11, 14–16, 24], which is deposited into the biomineralization compartment where crystallization occurs [15, 21, 25, 26].

The regulation of endocytosis and vesicular transport between membrane-bound cellular compartments were intensively studied in other systems [27, 28]. The observed size of the pocket and calcium vesicles in the sea urchin embryo (~1μm) suggests that this endocytosis process is most likely macropinocytosis (“cell drinking”), and not receptor/clathrin mediated endocytosis (upper limit of ~200nm) [27] or caveolin mediated (upper limit of ~80nm) [29]. Macropinocytosis is an actin-dependent process that initiates from surface membrane ruffles that give rise to large endocytic vacuoles called macropinosomes [30]. The macropinosomes go through a maturation process that involves shrinking and coating with various membrane-bound proteins [31, 32]. Some of the membrane-bound proteins could be motor proteins like dynein or kinesin that actively transport vesicles along the microtubules in the cells [33, 34]. This mechanism could, in principle, be used in calcifying cells, to direct the mineral-bearing vesicles into the biomineralizing compartment. Alternatively, the vesicles can perform a diffusive motion that is constrained by the cellular organelles and affected by the dynamic remodeling of the cytoskeleton network within the cell [35–40]. This mode of vesicle diffusion within the cells is called “active diffusion” to distinguish it from thermal diffusion [35–40]. Deciphering the mode of motion of the calcium vesicles within the skeletogenic cells and specifically, distinguishing between motor guided motion and diffusion, is essential for understanding the cellular regulation of mineral transport into the biomineralization compartment.

Previous studies of calcium vesicles in sea urchin embryos indicate that the calcium is processed differently in the calcifying skeletogenic cells compared to the non-calcifying cells [11, 14]. To study the calcium processing in the cells of sea urchin embryos, two dyes that can only enter the cells through endocytosis were added to the sea water: dextran-red and calcein dye, that marks calcium [11]. While most of the ectoderm vesicles were stained evenly with the two dyes, about a third of the skeletogenic vesicles were stained only in calcein and lack alexa-dextran. Furthermore, a recent study identified hundreds of vesicles where the calcium concentration was extremely high (1-15M) in the skeletogenic cells, while such vesicles were either very rare or completely absent in non-skeletogenic cells [14]. This indicates that calcium vesicles in the skeletogenic cells are biologically processed to eliminate the sea water and increase the calcium concentration, while in the non-skeletogenic cells this processing does not occur. Comparing vesicles behaviors, the skeletogenic and non-skeletogenic cells, could reveal common and distinct vesicle properties and illuminate shared vs. biomineralization specific regulatory mechanisms.

The molecular regulation of sea urchin skeletogenesis has been intensively studied resulting in a state-of-the-art model of the gene regulatory network (GRN) that controls skeletogenic cell fate specification [18, 19, 41–43]. A key control gene in this GRN encodes the Vascular Endothelial Growth Factor Receptor (VEGFR) [18, 42–45] that in vertebrates regulates the formation of blood vessels (vascularization) and the sprouting of new blood vessels from existing ones (angiogenesis) [46]. VEGFR is exclusively expressed in the sea urchin skeletogenic cells, and its inhibition by either genetic manipulation or using the VEGFR specific inhibitor, axitinib, distorts skeletogenic cell migration and completely blocks spicule formation [18, 42, 44]. We previously studied the role of sea urchin VEGF signaling in calcium vesicle accumulation using calcein labeling in live embryos visualized by confocal microscopy [18]. The number of calcium vesicles in the skeletogenic cells is unaffected by VEGFR inhibition until the time of spicule formation in normal embryos. When the spicules form in normal embryos the number of vesicles in the skeletogenic cells is higher under VEGFR inhibition compared to normal embryos. Possibly, calcium vesicles accumulate in higher numbers in the skeletogenic cells when VEGF signaling is inactive since the biomineralization compartment doesn’t form and vesicles are not deposited. Yet, these measurements were based on two dimensional images and lack the volumetric information and temporal resolution required for assessing calcium vesicle volume and kinetics in the 3-dimensional (3D) cellular environment. Lattice light-sheet microscopy (LLSM) is a promising technique for the measurement and assessment of the 3D vesicle dynamics in the cells of live sea urchin embryos [47].

LLSM is a recent improvement in 3D fluorescence imaging [47]. Light-sheet microscopy (LSM) excites a sample using an excitation objective that is perpendicular to the imaging objective. Lattice light-sheet techniques extend LSM by using a lattice beam to excite the sample rather than a Gaussian beam. The lattice beam improves the spatial resolution of the excitation, as well as reduces the intensity of excitation on the sample, while avoiding the photobleaching problems that are common with confocal and standard LSM. In addition, the reduced excitation of the LLSM avoids harming the cells of the embryo allowing for the long-term visualization of calcium dynamics.

The power of LLSM to produce fast high-resolution 3D imaging also presents new challenges for data analysis. Full 3D volumetric visualization provides a comprehensive view of the data, but it also requires a deep understanding of the image statistics in order to properly highlight structures of interest. Similar to looking through fog, image noise in 3D volumetric data visualization can obscure the structure unless the visualization parameters are tuned for each dataset. Manually quantifying cellular and sub-cellular dynamics in tens of 3D movies, each having hundreds of frames, is error-prone and in most cases, practically infeasible.

Automated analysis approaches are highly suitable for handling large 3D LLSM datasets. Quantification of cellular dynamics begins with automatic identification of individual structures of interest in each frame (often referred to as segmentation). After segmentation, a tracking algorithm links the segmented structures frame-to-frame to maintain the “identity” for each object over time. Once tracking is complete, size and motion dynamics can be measured for each object and statistical comparisons can be made. Several software tools have been developed focused on automatic 3D biological image analysis [48–50]. The LEVER package is extensible and supports visualization of the tracking along with the raw image data for validation of the automated results. LEVER has previously been used for both 2D and 3D cellular lineage analysis. As part of this work, we extended LEVER to support identification and tracking of subcellular structures such as calcium vesicles. In order to support this functionality, we integrated a new 3D vesicle segmentation approach, as well as improved the 3D visualization and tracking algorithms to better support subcellular dynamics analysis [51–53].

Here we used LLSM to measure the 3D dynamics of calcium vesicles in the calcifying skeletogenic cells and the non-calcifying ectodermal cells, in control and VEGFR inhibited sea urchin embryos at the gastrula stage. We extended the LEVER software platform to support automatic identification and tracking of calcium vesicles in 3D and to generate interactive volumetric 3D visualization and movie capture. Our studies indicate that calcium vesicles perform an active diffusion motion in both the skeletogenic and ectodermal cells, but vesicle speed and diffusion length are significantly different between these two cells types, possibly due to their distinct mechanical properties. To test this, we quantified the distribution of actin filaments and active myosin II and observed significant differences between the skeletogenic and ectodermal cells, in both normal and VEGFR inhibited embryos. Our studies reveal the typical volume and velocity scales and the characteristic motion of calcium vesicles in the sea urchin embryonic cells and point toward the cytoskeleton remodeling mechanisms as major effectors of this motion.

## Results

### Detecting cellular and calcium vesicle dynamics in LLSM

In order to visualize 3D calcium vesicle dynamics in live sea urchin embryos for long periods of time, we used the LLSM at the advanced imaging center at the Janelia research campus [47]. This LLSM setup allows for high temporal resolution of ~40ms per slice and about ~2 sec for 3D embryonic volume of ~40μm^3^ [47, 54]. We used green calcein to stain the calcium ions [15] and FM4-64 to mark cell membranes in live sea urchin embryos of the species, *Lytechinus variegatus* (*L. variegatus*) at the gastrula stage (Fig. 1, [18]). To study the effect of VEGF signaling we treated sea urchin embryos with the VEGFR inhibitor, axitinib. Axitinib is a specific inhibitor of human VEGFR2 that binds to the kinase domain that is highly conserved between human and sea urchin; specifically, the six amino-acids to which Axitinib binds are conserved between human and sea urchin VEGFR [18]. Axitinib treatment results in similar skeletogenic phenotypes to those observed in genetic perturbations of the VEGF gene in *L. variegatus* and other species [18, 42] and particularly, it results in complete skeletal loss [18, 42]. Axitinib is dissolved in DMSO and our control embryos were therefore cultured in a similar concentration of DMSO like the axitinib embryos, see Methods for experimental details.

**Figure 1.**
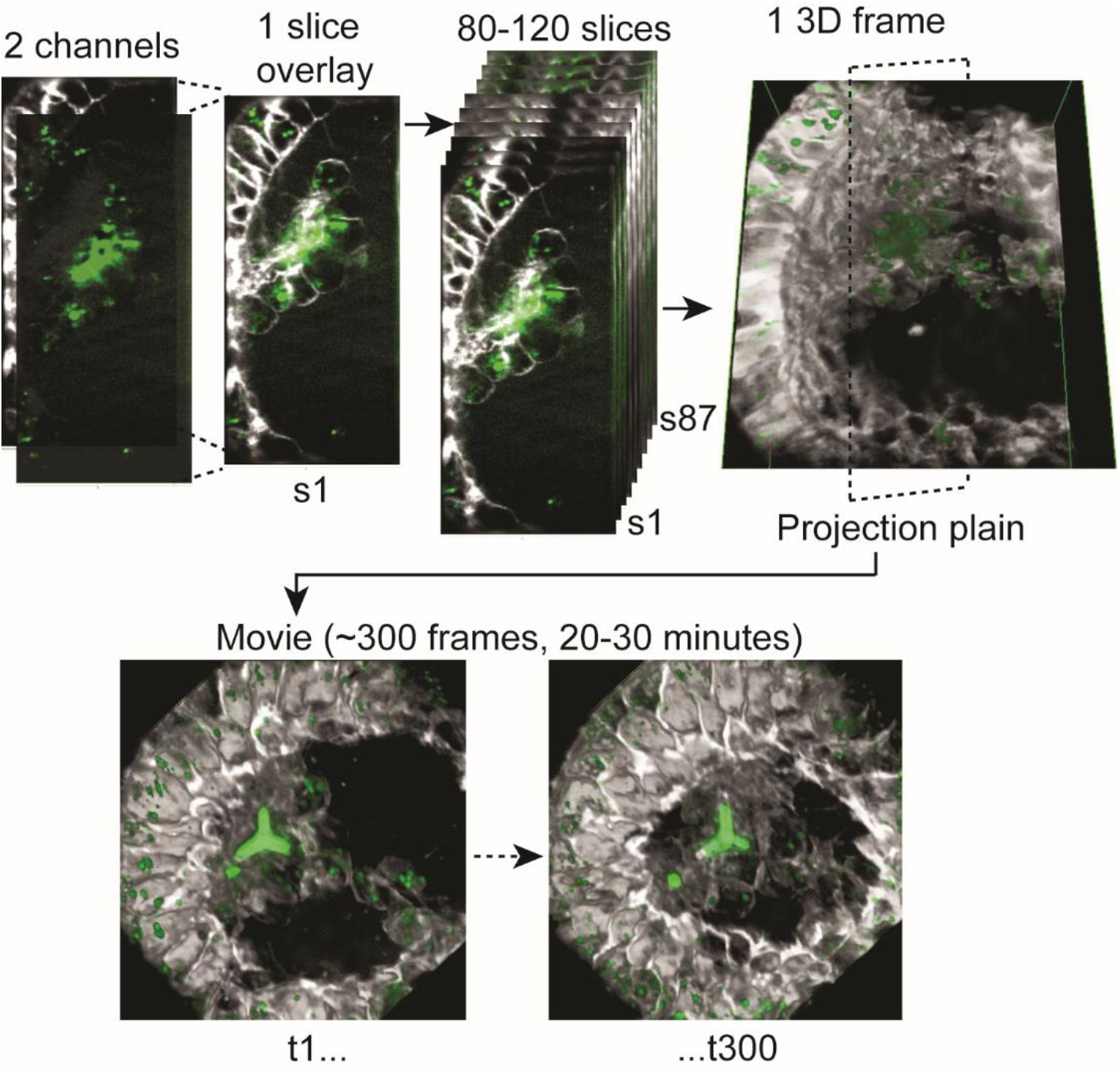
4D lattice light sheet image processing. Each slice is composed from two fluorescent channels, the calcein (green) that marks the calcium and the FM4-64 (white) that marks cell membranes. Each time points (frame) is composed of 80-120 slices that are reconstructed to form a 3D image of the volume of the embryos, as demonstrated in movie S1. We then project the 3D image into a selected plane and generate movies showing 200-400 consecutive frames spanning about 20-30 minutes.

Before we further describe our experiments and observations, some methodological limitations must be mentioned. The embryos were grown in calcein until early gastrula stage and then calcein was washed about two hours or more before we could image vesicle dynamics. This time interval after the wash was necessary to eliminate the calcein stain from the cell cytoplasm and blastocoelar fluid so the individual calcein stained-vesicles would be distinct from the background. However, as the blastocoelar fluid was not stained with calcein during the imaging, we were unable to detect live-pinocytosis as well as unstained calcium vesicles that were apparently deposited into the spicules. To immobilize the embryos and enable live-imaging for long period of time (>30min), we immersed the embryos in low melting agarose. In this condition the spicule elongation is significantly slower than in free swimming embryos. The agarose stiffness is about 35-50kP [55] which is much more rigid than the sea water where the sea urchins naturally grow. This, and possibly the lack of movement, could underlie the slow growth of the spicule under these conditions.

### 4D rendering of the LLSM data reveals rich cellular behaviors and vesicular dynamics

To visualize the 3D structure and motion of the cells and the vesicles in the embryos throughout time, we reconstructed a 3D model from the individual slices and then connected these images over time to produce time-lapse sequences (Fig. 1). The image of each slice is made up of two-channels, calcein (green) and FM4-64 (white). Each time point (frame) is built from 80 to 120 image-slices that are reconstructed into a 3D volume that spans approximately 40μm^3^ of the sea urchin embryo as demonstrated in Fig. 1 and in Movie S1. We then project the 3D volume onto a viewing plane to enable 2D visualization of the time lapses (movies, Fig. 1). We recorded nine control embryos from five different sets of parents and eleven VEGFR inhibited embryos from four different sets of parents. Details of all these time-lapses are provided in Dataset S1 where in each date a separate set of parents was recorded. We present here six representative movies, three of control embryos and three of VEGFR inhibited embryos, that include 200-400 frames separated by time intervals of 3.26-6.12 sec, spanning about 20-30 minutes of cellular and vesicular motion (Figs. 2, 3 and Movies S2-7).

**Figure 2.**
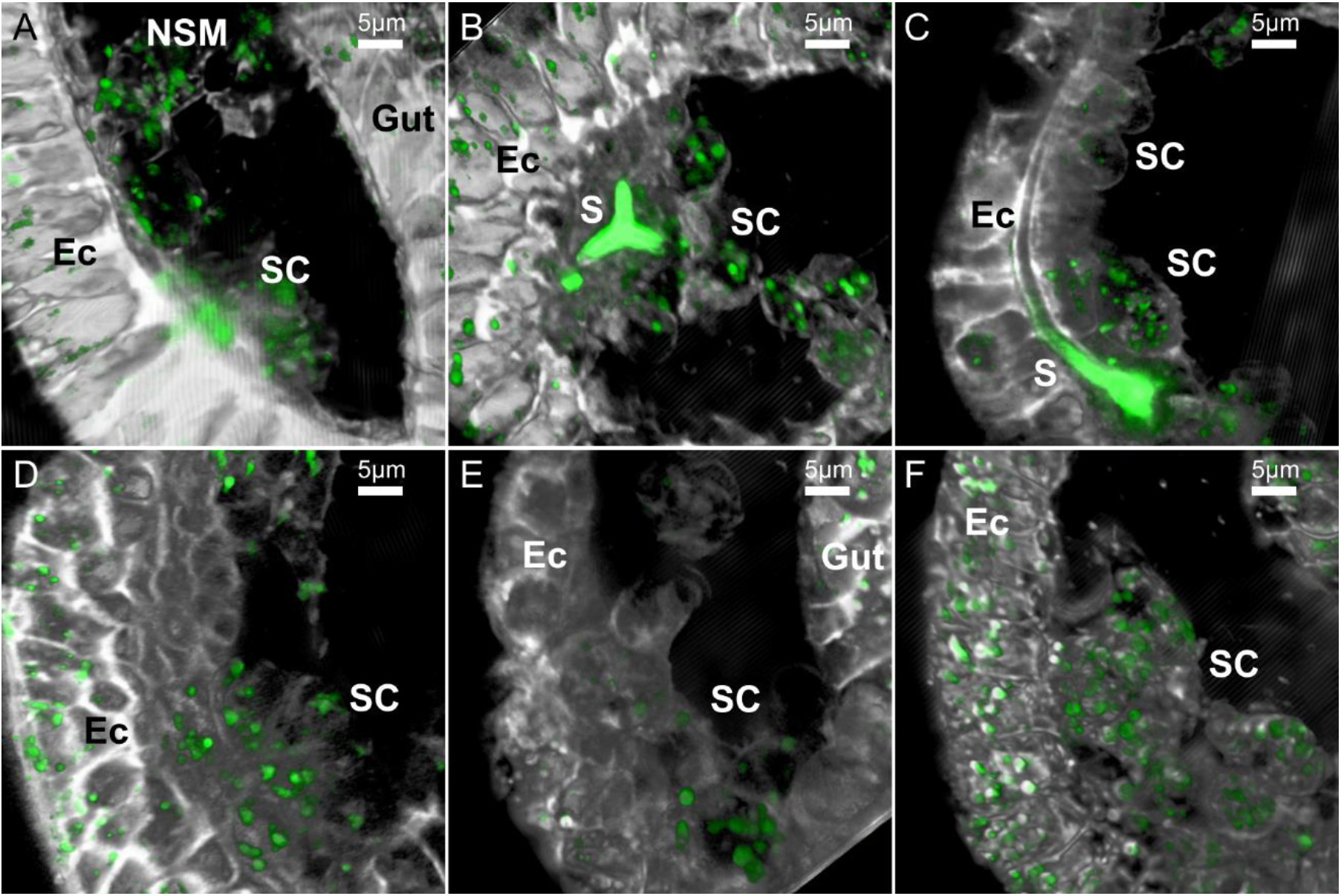
Examples of the LLSM 3D images of sea urchin embryos in different developmental stages and treatments. These representative images are 2D projections of the 3D rendered frames of selected datasets, as demonstrated in Fig. 1. Calcein staining is marked in green and the FM4-64 membrane staining is marked in white. (A-C), control embryos (DMSO) at the early gastrula stage before spicule formation (A), just after the tri-radiate spicule forms (B) and when the spicule is elongated (C). (D-F) Representative embryos at gastrula stage treated with VEGFR inhibitor, axitinib, do not have spicules. Scale bars are 5μm. 3D movies showing the first 100-200 frames of each of the dataset presented in this figure are provided as movies S2-7. Ec – ectoderm, SC – skeletogenic cells, NSM-non-skeletogenic mesoderm, S – spicule.

**Figure 3.**
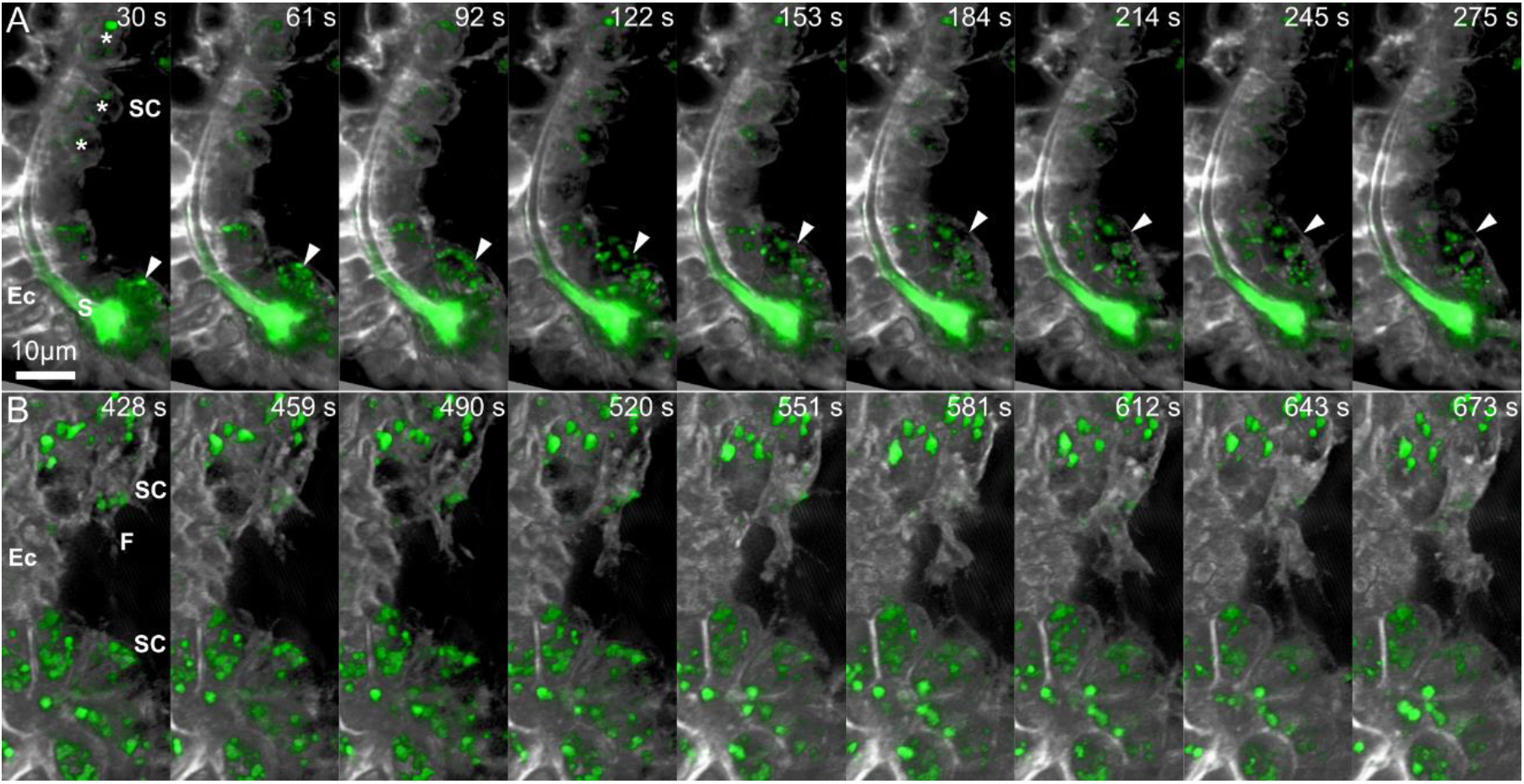
Sequences of time-lapse images demonstrating cellular dynamics in control and under VEGFR inhibition. (A) Time-lapse images of control embryo (Movie S4) showing the rapid movement of a free mesenchymal cell (marked in arrowhead) compared to the stable position of the skeletogenic cells that are in direct contact with the ectoderm (asterisks). (B) Time lapse images of an embryo grown under VEGFR inhibition demonstrating the active filopodia extension and fusion between two skeletogenic cell clusters. Relative time from the beginning of the movie is shown in seconds at the top of each frame. Scale bar is 10μm. Ec – ectoderm, S – spicule, SC – skeletogenic cell, F - Filopodia.

The rendered movies demonstrate the highly dynamic cellular and vesicular motion in the two experimental conditions. In both conditions, the skeletogenic cells are in close contact with the ectodermal layer. In normal embryos the skeletogenic cells are in close vicinity to the developing spicule (Fig. 2B,C, 3A) and in both conditions they form clusters and rapidly extend filopodia that contact and fuse between cells [42] (Figs. 2 and 3). We also detect some non-skeletogenic mesenchymal cells that move around individually and are not in direct contact with the ectoderm (Fig. 2A, 3A, Movies S2, S4). The formation of skeletogenic cell clusters that are bound to the ectoderm in both control and VEGFR inhibition suggests that VEGF-independent ectodermal cues are responsible for the observed skeletogenic adhesion [56].

### Vesicle volume is larger in the skeletogenic cells vs. the ectodermal cells and increases under VEGFR inhibition

Next, we wanted to study the volume of calcium vesicles within the different embryonic tissues in normal and VEGFR inhibited embryos. To automatically quantify vesicle volume, we identified the 3D structure and volume of each vesicle in all frames using the newly developed image processing filters as part of the segmentation algorithm discussed in the methods section (demonstrated in Fig. 4A, B, and Movie S8). After automated identification of each vesicle, the size of each vesicle can be measured based on the volume of the pixels identified. To differentiate between the skeletogenic cells and the other embryonic territories we manually identified the boundaries of the ectoderm and of the endoderm regions in the first frame of each movie and used this frame for volume statistics (Fig. 4C). As the endoderm region was apparent only in few of the movies, we focused on characterizing the sizes of vesicles in the ectoderm and the skeletogenic cells. We measured vesicle volume in the nine control embryos and the eleven VEGFR inhibited embryos. The number of vesicles that were measured in the skeletogenic cells and in the ectoderm for each embryo is provided in Dataset S1. In some of the movies the mesodermal region contains a few non-skeletogenic mesodermal cells, but since the movies were focused on clusters of skeletogenic cells, the contribution of the non-skeletogenic cells to our analyses is minor.

**Figure 4.**
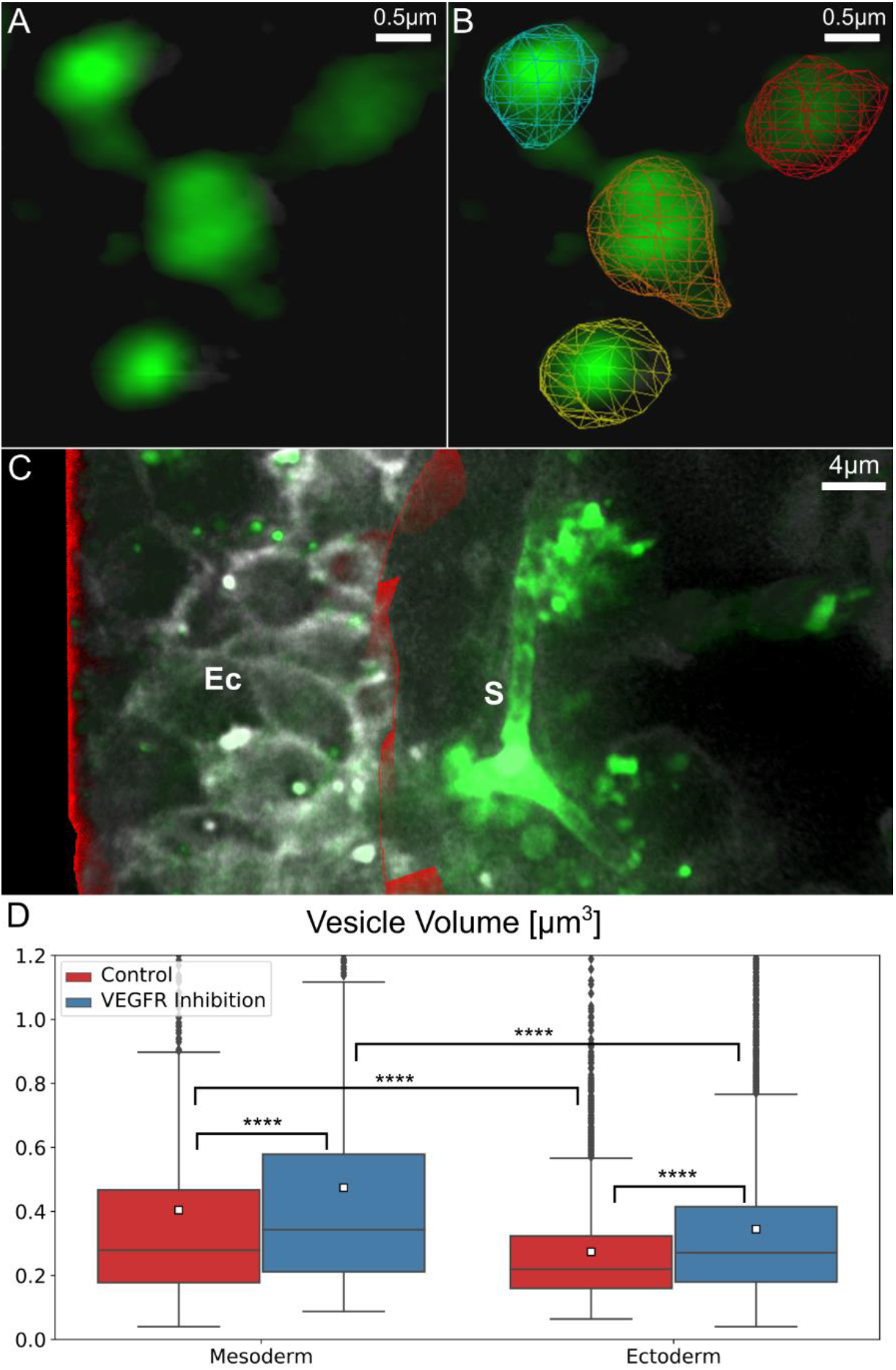
Vesicle volume is larger in the skeletogenic cells compared to the ectodermal cells and is significantly larger under VEGFR inhibition. (A-C) An example for the image processing involved in the quantification of vesicle volume in the ectodermal vs. skeletogenic embryonic domains. (A) Raw image rendering of calcium vesicle detection. (B) Demonstration of the automated vesicle detection (segmentation) overlaid on the image in (A). (C) Manual identification of ectodermal region in red, rendered along with raw image frame. (D) Comparison of vesicle sizes in ectodermal and skeletogenic cells in control and VEGFR inhibition. The total number of vesicle measured in the control skeletogenic cells: 815, ectodermal cells: 3719; VEGFR inhibition skeletogenic cells: 1530, ectodermal cells: 8621. Each box plot shows the average (white square), median (middle line), the first and the third quartiles (the 25^th^ and 75^th^ percentiles, edges of boxes) and outliers (black dots). Vesicle volume is significantly higher in the skeletogenic cells compared to the ectodermal cells and VEGFR inhibition significantly increases vesicle volume in both cell types. (Dunn-Sidak test, p<0.0001, exact p-values are given in Dataset S1).

Our measurements show that in control embryos, the average volume of calcein stained vesicles in the skeletogenic cells is ~0.4μm^3^ while in the ectodermal cells it is significantly smaller, ~0.27μm^3^ (Fig. 4D, Dataset S1). Under VEGFR inhibition, the average volume of calcein stained vesicles is higher compared to normal embryos both in the skeletogenic cells and in the ectodermal cells (Fig. 4D, Dataset S1). The average volume of the calcein stained vesicles under VEGFR inhibition is ~0.47μm^3^ in the skeletogenic cells and ~0.35μm^3^ in the ectodermal cells, an increase of more than 17% in vesicle volume in both tissues. We previously observed an increase in the number of vesicles in the skeletogenic cells under VEGFR inhibition at the time when the spicule forms in normal embryos [18]. The increase in vesicle number and volume might result from the higher level of calcium present in the blastocoel when VEGF signaling is inactive, since calcium is not sequestered into the spicules and accumulates in the blastocoel. Hence, when the cells perform endocytosis they uptake higher concentrations of calcium which leads to the observed increased calcium content in both the ectoderm and the skeletogenic mesodermal cells.

### Vesicle speed is slower in the skeletogenic cells compared to the ectodermal cells and the directionality is similar in the two tissues

We wanted to quantify vesicle dynamics in the cells of the sea urchin embryo and study the effect of VEGFR inhibition on vesicle motion. To do that we updated the tracking algorithm we developed previously [53, 57] to improve long-term tracking performance (see the methods section for details). Tracking maintains the identity of each vesicle over time, allowing instantaneous speed and velocity measurements, frame to frame, for each individual vesicle (demonstrated in Movie S9). Throughout this manuscript we use the term “**speed**” to describe the size of the velocity and the term “**velocity**” to describe the vector-velocity that includes the information of both the size and direction. An example **instantaneous speed** measurement is shown in Figure 5A, the magenta line represents the motion of the vesicle from the previous frame. The vesicle instantaneous speed is the length of the line (distance traveled in a single frame) divided by the time between frames (here, ~6 seconds). An important constraint on the effectiveness of tracking algorithms is the temporal resolution of the imaging relative to the speed of tracked objects. In the case of the calcium vesicles, we discovered that too much motion occurred between frames for effective tracking if the temporal intervals were 15 seconds or more between frames. We therefore only applied the tracking algorithm and motion analyses to movies with temporal resolution that is faster than 15 seconds between frames. To have equivalent time intervals in the two experimental conditions, we analyzed five control data sets and four VEGFR inhibition datasets, where the time intervals between frames is about 6 seconds (see Dataset S1 for the number of vesicles analyzed in each territory in each movie that was included in this analysis).

**Figure 5.**
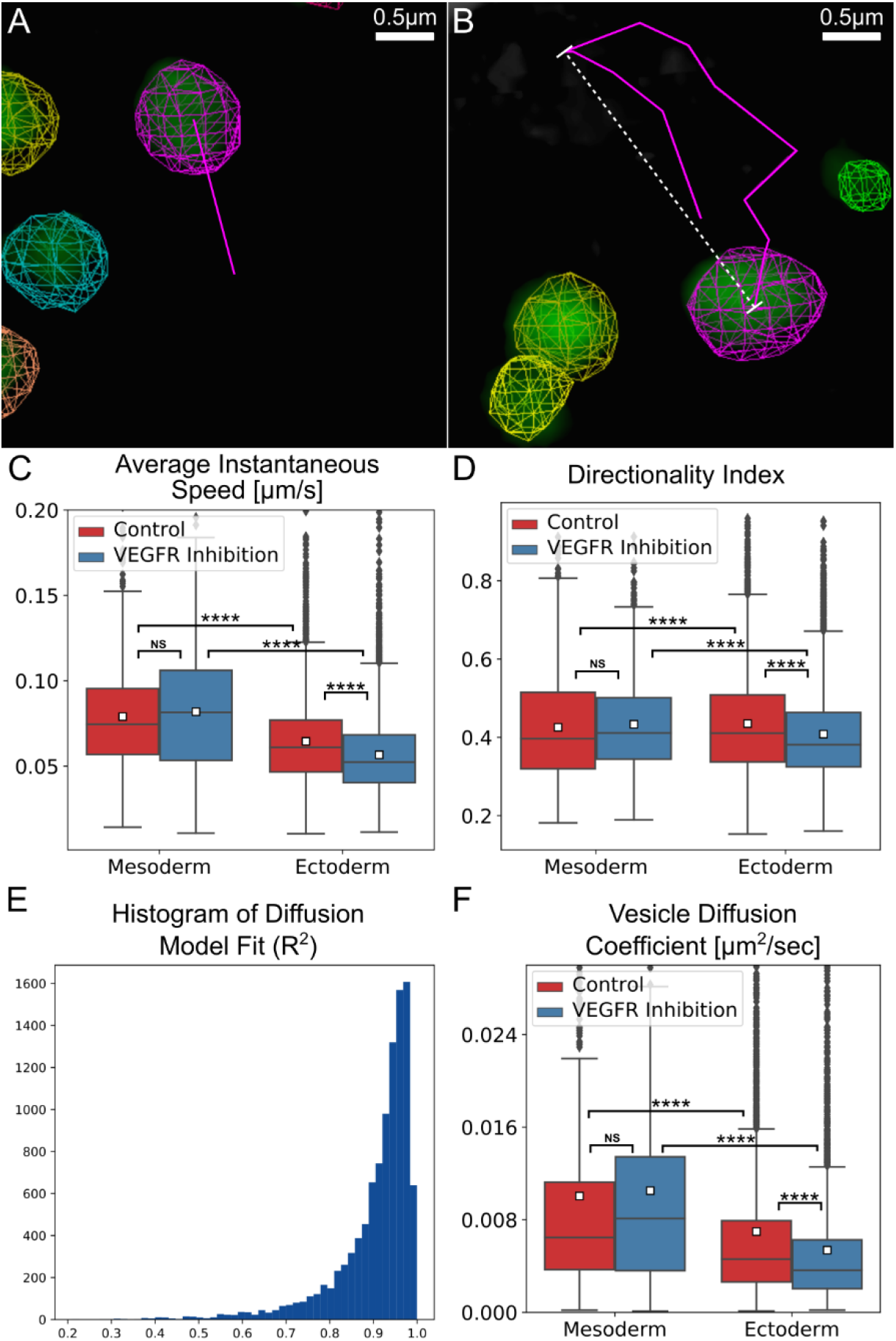
Vesicle tracking reveals an active diffusion motion with higher diffusion coefficient and speed in the skeletogenic cells compared to the ectodermal cells. (A, B) Examples for the automated tracking used to quantify vesicle kinetics. (A) Instantaneous speed indicates the distance traveled between sequential frames divided by the time interval between the frame. The magenta line demonstrates this distance for the magenta labeled vesicle. (B) Directionality index is the ratio of maximal displacement (white line) over the total distance traveled (magenta line) within a one-minute time interval. A representative 3D movie of tracking session is provided in movie S7. (C-E) Comparison of vesicle motion statistics between control and VEGFR inhibited embryos. The total number of vesicle measured in the control skeletogenic cells: 825, ectodermal cells: 4193; VEGFR inhibition skeletogenic cells: 884, ectodermal cells: 4508. Each box plot shows the average (white square), median (middle line), the first and the third quartiles (the 25^th^ and 75^th^ percentiles, edges of boxes) and outliers (black dots). (C) Vesicle instantaneous speed. (D) Directionality index. (E) Vesicle diffusion coefficient. (Dunn-Sidak test, p<0.0001, exact p-values are given in Dataset S1). (F) Histogram of Diffusion model fit, 90% of vesicle tracks are well modeled by the standard diffusion model (R^2^>0.8).

Our measurements show a distinct difference in the instantaneous speed between the skeletogenic and the ectodermal cells. The average instantaneous speed per track of the vesicles in the skeletogenic cells is about 0.079μm/sec in control embryos and it is not affected significantly by VEGFR inhibition (Fig. 5C, Dataset S1 and methods). A significant difference is observed between the vesicle speed in the skeletogenic cells and in the ectoderm, where the average instantaneous speed per track is 0.065μm/sec in control embryos. Vesicles speed in the ectodermal cells reduces to ~0.057μm/sec under VEGFR inhibition. Thus, vesicle volume and speed are larger in the skeletogenic cells compared to the ectodermal cells in both control and VEGFR inhibition (Fig. 4D, 5C, Dataset S1). Despite these clear trends, there is no correlation between vesicle volume and speed throughout the experimental conditions (Fig. S1A, B). The lack of correlation implies that the change of vesicle volume is not related to the change in vesicle motion and is due to different regulation of these quantities in the embryonic cells.

To further characterize vesicle motion, we quantified the directionality of vesicle velocity by measuring the **directionality index**. The directionality index is the ratio between the maximal displacement and the total length of the vesicle track during a time window of one minute (Fig. 5B) [58, 59]. Values close to 1, reflect linear (directed) motion and values close to 0, reflect random motion. The average directionality index in both the skeletogenic and ectodermal cells is about 0.43 and under VEGFR inhibition it doesn’t change significantly in the skeletogenic cells and shows a minor change in the ectodermal cells (Fig. 5D, Dataset S1). This value is similar to the directionality index measured for endocytic vesicles that are guided by molecular motors along microtubules in astrocyte cultures [58, 59]. However, the typical speed of motor-guided vesicles in these cultures and in other systems is much faster (~0.4-0.5μm/sec) [58–60] compared to our measurements (~0.06-0.08μm/sec). Additionally, the motor-guided movement of vesicles along microtubules was shown to be in bursts of directed motion that last ~6-7 seconds followed by a pause for a similar time interval [36, 60]. If that was the case for the calcium vesicles, we would have expected to see two peaks of instantaneous speed, a fast-speed peak corresponding to the directed motion and a slow-speed peak corresponding to the pause. However, the distribution of measured instantaneous speed shows only a single peak that corresponds to the relatively slow movement of the vesicles reported above (Fig. S2). The slow instantaneous speed and the lack of two modes of motion suggest that the vesicle motion is not guided by molecular motors along the microtubule network.

### The calcium vesicles perform a diffusion motion in the skeletogenic cells and the ectoderm

In other systems, vesicles were shown to experience a diffusive motion that is not thermal in nature but results from the dynamic remodeling of the cytoskeletal network and the ubiquitous activity of motor proteins that affect every moving object within the cell cytoplasm [36–40]. This mode of diffusion is called “active diffusion” and is characterized by a larger amplitude (step size) compared to the amplitude of thermally-induced diffusion. Additionally, the diffusion coefficient for active diffusion is independent of particle size, unlike thermal diffusion where the diffusion coefficient decreases with increasing particle size [37]. To test whether the calcium vesicles experience an active diffusion motion we fit a standard diffusion motion model to each track. We plot the mean-square displacement, 〈Δ*x*^2^〉, as a function of time, Δ*t*, and use a linear fit to measure the **diffusion coefficient** [61], ***D***, 〈Δ*x*^2^〉 = *D*Δ*t*, for each track (Fig. S3). Mean-square displacement is measured by taking the squared sum of path-lengths (*e.g.* the lengths of magenta line segments in Fig. 5B) for each vesicle track. Δ*t* is the time interval between two consecutive frames, which in our data is 5.6-6.12 seconds (exemplified in Fig. S3). Most of the data fit well within this model, that is, *R*^2^ ≥ 0.8 for 90% of the tracked vesicles (Fig. 5E). The average diffusion coefficient is D~0.01*μm*^2^/*sec* in the skeletogenic cells and is not affected by VEGFR inhibition (Fig. 5F, Dataset S1). In the ectoderm the diffusion coefficient is smaller, D~0.007*μm*^2^/*sec* and it decreases to ~0.005*μm*^2^/*sec* under VEGFR inhibition. The lower diffusion coefficient in the ectodermal cells is in agreement with the lower instantaneous speed measured for the ectodermal vesicles compared to the skeletogenic cells (Fig. 5C). The diffusion coefficient does not correlate with vesicle size (Fig. S4), which supports an active diffusion motion. Thus, our analyses indicate that calcium vesicles perform a diffusion motion that has the signature of an active diffusion, in both the skeletogenic and ectodermal cells, with higher diffusion coefficient in the skeletogenic cells.

### Vesicle motion is not directed toward the spicule but the vesicle speed slows down close to the spicule

We wanted to investigate vesicle deposition in normal embryos and study vesicle motion near the spicule in this condition. Vesicle-membrane fusion is a very fast process that occurs within ~100 milliseconds [62], and therefore our temporal resolution did not allow us to detect such events. However, vesicle content deposition can last several minutes, as was shown for the vesicles that carry adhesive glycoproteins secreted into the lumen of the drosophila salivary gland [63, 64]. To see if we could detect such processes, we studied the directionality and speed of vesicle motion relative to the spicule and tried to infer whether the vesicles are trafficked toward the spicule and if they slow down near the spicule vicinity.

To measure vesicle motion relative to the spicule we manually marked the spicule and measured vesicle velocity toward the spicule at increasing distances from the spicule for three movies where the spicule was evident (Fig. 6A, B). The average velocity is around zero indicating random motion toward and away from the spicule, which suggests that vesicle motion is not directed towards the spicule; in other words, the spicule does not attract vesicle movement towards it. However, the average vesicle speed is significantly lower near the spicules (Fig. 6C). The vesicle speed is ~0.05μm/sec at distances of 1-2μm from the spicule, while at distances ≥7μm it increases to ~0.08μm/sec, which is the average instantaneous speed in the skeletogenic cells (Fig. 6C). Overall, vesicle velocity is not directed toward the spicules but the vesicles significantly slow down near the spicule, possibly, as they bind to it and deposit their content.

**Figure 6.**
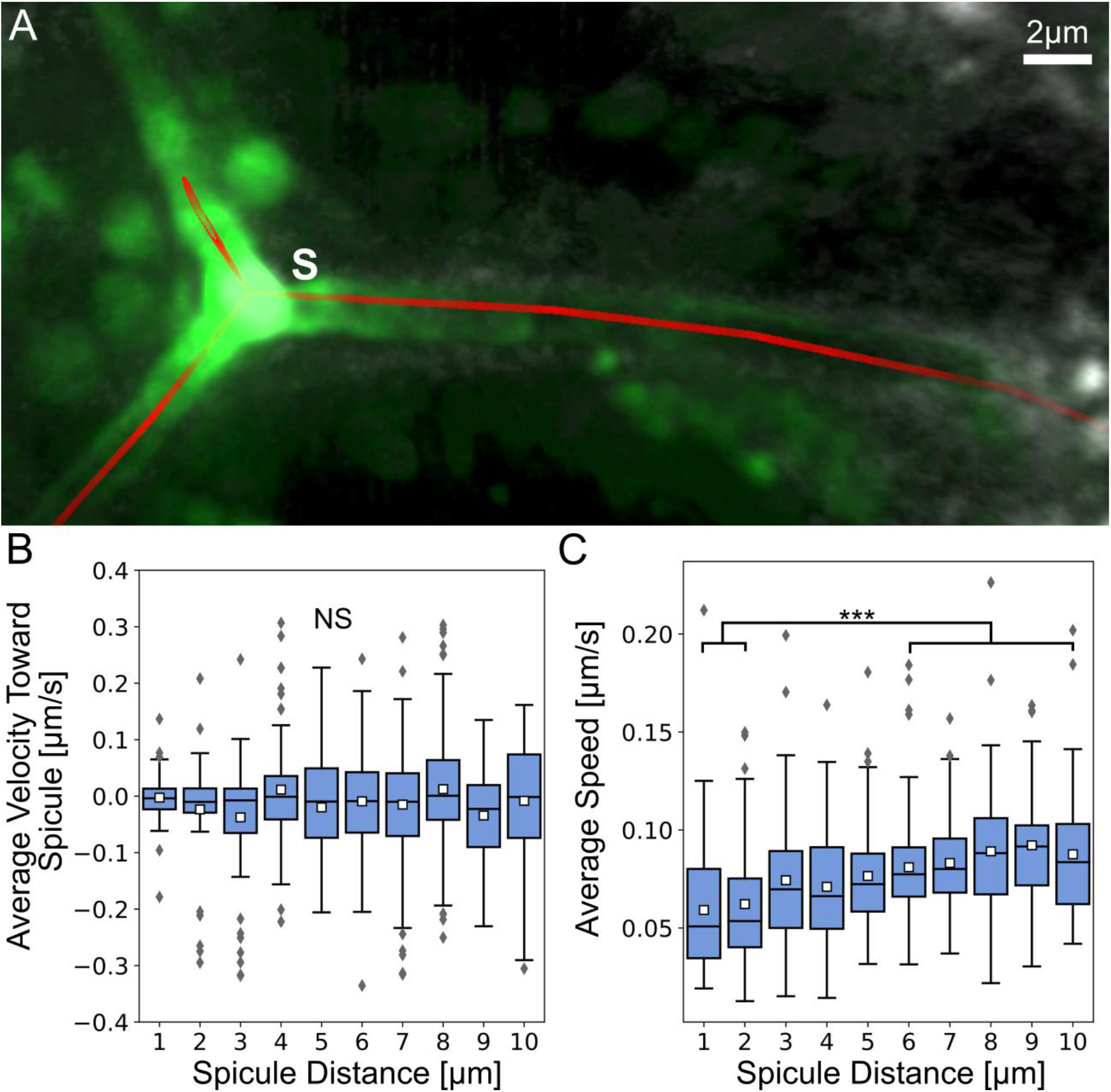
Vesicle velocity is not directed toward the spicule but vesicle speed is lower near the spicule. (A) An example for the manually identified spicule centerline shown in red with raw image data. (B) Average instantaneous velocity (μm/sec) toward the spicule relative to the average distance from spicule. Each box plot shows the average (white square) median (middle line), the first and the third quartiles (the 25^th^ and 75^th^ percentiles, edges of boxes) and outliers (gray diamonds). (C) Average instantaneous vesicle speed (μm/sec) at increasing average distances from the spicule (μm). The speed at distances 1-2μm are significantly lower than at distances >6μm (Dunn-Sidak test, p<0.001, exact p-values are given in Dataset S1), with n=803 vesicles.

### The spicule is enriched with f-actin but the skeletogenic cells show lower strength and activity of the actomyosin network compared to ectodermal cells

Our analyses, in the light of previous studies, point toward cytoskeleton remodeling as an important effector of vesicle motion and deposition. Active diffusion depends on the dynamics of cytoskeleton remodeling and is affected by the strength and activity of the actomyosin network [37–40]. Furthermore, vesicle deposition was shown to be regulated by a network of cytoskeleton remodeling proteins that controls f-actin assembly and myosin contractility [64]. We therefore wanted to study the strength and activity of the actomyosin networks in the skeletogenic and ectodermal cells in normal and VEGFR inhibited sea urchin embryos. To identify the skeletogenic cells, we used the marker, 6a9, that binds to the skeletogenic specific cell surface glycoprotein, mspl30, [65]. We used red phalloidin to identify actin filaments (f-actin) and the Anti-Myosin light chain (MyoIIP) antibody to detect activated myosin in the embryos of the species, *Paracentrotus lividus* (*P. lividus*).

The spicule is enriched with f-actin but the skeletogenic cells show lower f-actin signal compared to the ectodermal cells (Fig. 7). f-actin is enriched in the apical side of the ectodermal cells (arrowhead, Fig. 7A), in agreement with previous studies [66]. This enrichment is observed in both control and VEGFR inhibited embryos (Fig. 7A, D). Strong signal of f-actin is clearly detected around the spicule of normal embryos, indicating that an actomyosin network is assembled near the spicule membranes (Fig. 7A, C), in agreement with previous studies in sea urchin skeletogenic cells cultures [67]. f-actin is also observed near the cell membranes in normal and VEGFR inhibited embryos (Fig. 7A, C, D, F). To quantify the density of f-actin in the different cells and experimental conditions we measured the number of red pixels for area in the skeletogenic cells and in the ectodermal cells (Fig. 7G, see methods for details). The f-actin signal is significantly stronger in the ectodermal cells compared to the skeletogenic cells and it is unaffected by VEGFR inhibition in both cell types (Fig. 7H). Thus, f-actin is assembled around the spicule membranes in normal embryos and f-actin is enriched in the epithelial ectodermal cells compared to the mesenchymal skeletogenic cells, in both normal and VEGFR inhibited embryos.

**Figure 7.**
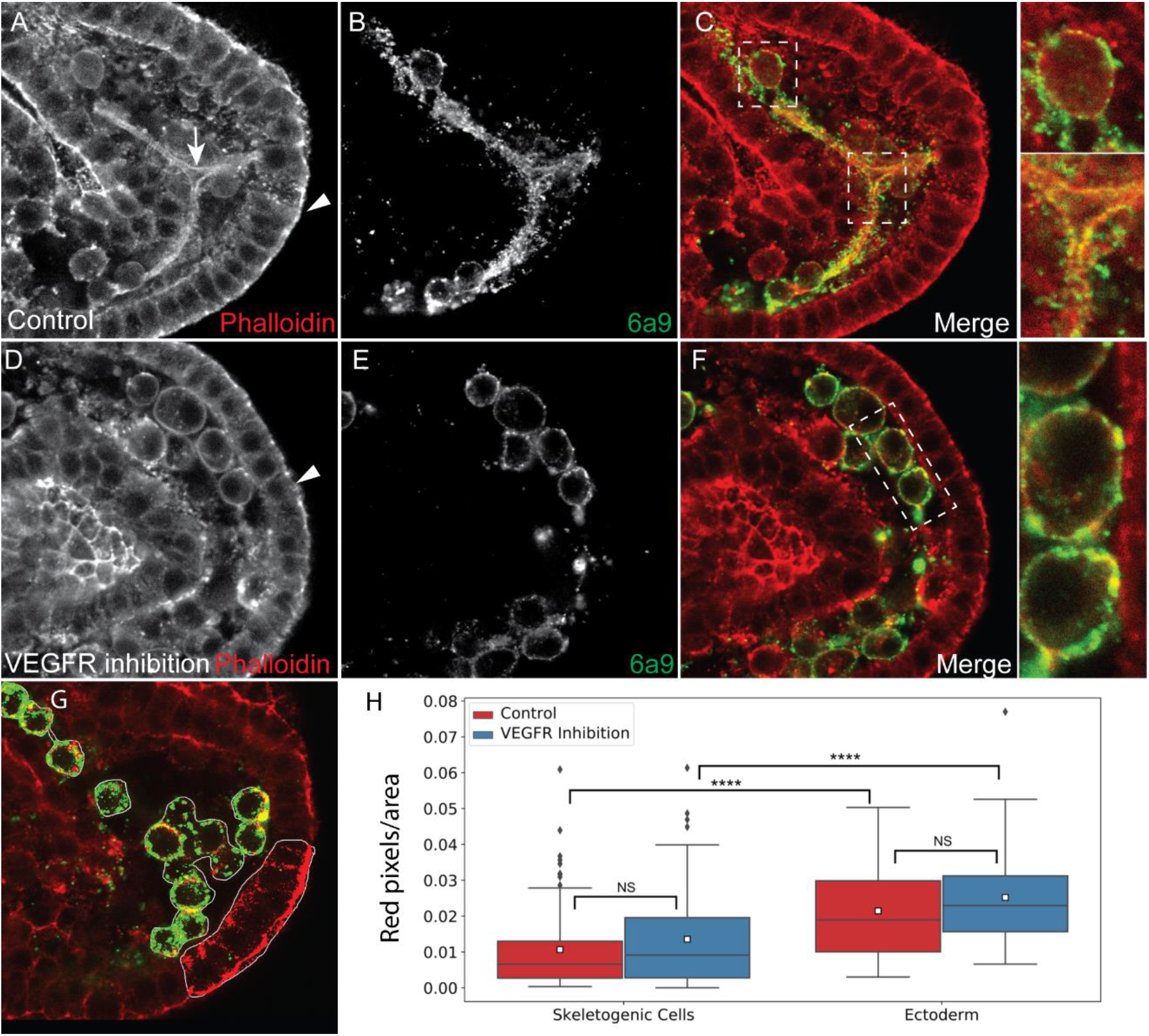
Actin filaments are detected around the spicule and are enriched in the ectodermal cells compared to the skeletogenec cells. (A-F) Representative images showing actin filaments in normal embryos (A-C) and VEGFR inhibited embryos (D-F). Phalloidin was used to stain f-actin (A and D) and 6a9 was used to mark the skeletogenic cells (B and E). Arrow in A marks the spicule, arrowheads in A and D mark the apical side of the ectodermal cells. In C and F we present the overlay of the phalloidin and the skeletogenic marker, with indicated sections enlarged on the right. (G-H) quantification of the phalloidion signal. The number of red pixels (phallodin signal) per marked area was measured (G). The phallodin signal is significantly higher in the ectodermal cells compared to the skeletogenic cells and it unaffected by VEGFR inhibition (Dunn-Sidak test, p<0.0001, exact p-values are given in Dataset S1). Based on 3 biological replicates where overall n=24 normal embryos and n=30 VEGFR inhibited embryos were studied.

The signal of active myosin is much stronger in the ectodermal cells compared to the skeletogenic cells supporting a higher activity of the actomyosin network in the ectodermal cells (Fig. 8). Active myosin is enriched in the apical side of the ectodermal cells in both normal and VEGFR inhibited embryos (arrowhead in Fig. 8A and D). Differently than f-actin, active myosin is not enriched around the spicules in normal embryos (Fig. 8A, C). To quantify the level of active myosin we measured the number of red pixels per area in the skeletogenic cells and in the ectodermal cells of normal and VEGFR inhibited embryos (Fig. 8G). Active myosin signal is significantly higher in the ectodermal cells compared to the skeletogenic cells and is it unaffected by VEGFR inhibition in both cell types (Fig. 8H).

**Figure 8.**
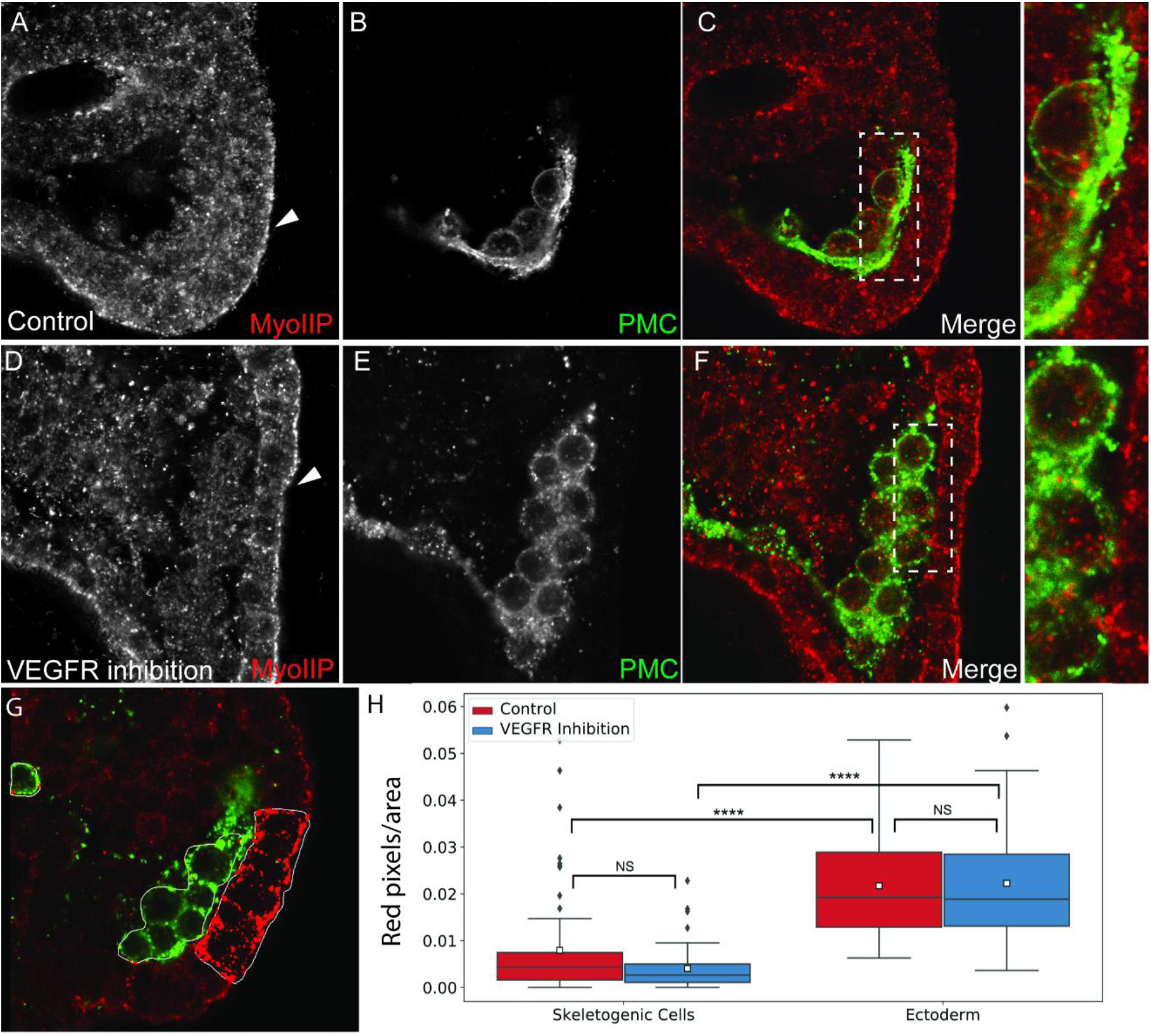
Active myosinII signal is enriched in the ectodermal cells compared to the skeletogenec cells. (A-F) Representative images showing active myosinII (MyoIIP) in normal embryos and VEGFR inhibited embryos. (A, D) active MyosinII in normal and VEGFT inhibited embryos, respectively. Arrowhead indicates the apical side of the ectodermal cells. (B, E) skeletogenic cells marker (6a9) in normal and VEGFR inhibited embryos, respectively. (C, F) overlay of the two markers, with indicated sections enlarged on the right. (G-H) quantification of the active myosinII signal. The number of red pixels (active myosinII) per marked area was measured (G). The active myosinII signal is significantly higher in the ectodermal cells compared to the skeletogenic cells and is unaffected by VEGFR inhibition (Dunn-Sidak test, p<0.0001, exact p-values are given in Dataset S1). Based on 3 biological replicates where overall n=27 normal embryos and n=30 VEGFR inhibited embryos were studied.

Overall, both f-actin and active myosin signals are significantly stronger in the ectodermal cells compared to the skeletogenic cells, suggesting a lower strength and activity of the actomyosin network in the skeletogenic cells. The lower strength and activity of the actomyosin network could underlie the higher vesicle speed and diffusion length observed in the skeletogenic cells, further supporting the active diffusion mode of vesicle motion.

## Discussion

A key requirement of biomineralizing organisms is the ability to accumulate minerals inside intracellular vesicles where the mineral is kept in an amorphous state until it is deposited into the biomineralization compartment [11, 13, 16]. Despite the key role of mineral bearing vesicles in the biological regulation of biomineralization, very little is known about their trafficking inside the biomineralizing cells and the regulation of their dynamics and deposition. Here we studied the cellular dynamics and the motion of calcium vesicles in calcifying and non-calcifying cells of normal sea urchin embryos and of embryos grown under VEGFR inhibition, where skeletogenesis is blocked. Our studies reveal differences in vesicle volume and speed between the non-calcifying ectodermal cells and the calcifying skeletogenic cells. Our data support an active diffusion motion of the vesicles in both cell types, with higher diffusion coefficients and lower strength and activity of the actomyosin network in the skeletogenic cells. The motion of the vesicles is not directed toward the spicule yet, they seem to slow down near it, as they possibly bind and deposit the mineral. The spicule is coated with f-actin, implying a possible role of the actomyosin network in supporting the spicule structure and regulating vesicle deposition. Below we discuss the possible molecular mechanisms that underlie these observations and the implications of our studies on the understanding the biological regulation of biomineralization and reproducing it in artificial systems.

Our measurements show that the average volume of calcein stained vesicles is larger in the skeletogenic cells compared to the ectodermal cells, in both normal and VEGFR inhibited embryos (Fig. 4D, Dataset S1). This difference might result from different processing of the vesicle content in these two cell populations. Indeed, previous studies indicated that calcium vesicles in the skeletogenic cells are biologically processed to eliminate the sea water and increase the calcium concentration, while in the non-skeletogenic cells this processing does not occur [11, 14]. Together, these findings indicate that the biological regulation of calcium vesicle content is distinct between the skeletogenic and the ectodermal cells which apparently leads to higher calcium vesicle volume in the skeletogenic calcifying cells. There is a multitude of evidence of a specific activation of genes that regulate 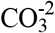 homeostasis in the skeletogenic cells, *e.g.,* the carbonic anhydrase like-7, Caral7 [18, 44] and the bicarbonate transporter, SCL4a10 [68]. However, further studies are required to identify the genes responsible for the specific regulation of Ca^+2^ ions in the vesicles of the skeletogenic cells.

VEGFR inhibition increases calcium vesicle volume in both the skeletogenic and ectodermal cells (Fig. 4D). Apparently, when the skeleton doesn’t form and the calcium is not sequestered into the spicule, calcium is accumulated in the sea urchin blastocoel and is taken by the cells through macropinocytosis. As VEGFR is only expressed in the skeletogenic cells, the increase of calcium vesicle volume in both tissues supports VEGF-independent macropinocytosis as a major source of calcium accumulation in all the cells of the embryo.

Calcium vesicles perform an active diffusion motion in all the cells of the embryo, as evident from their relatively slow speed, the good agreement of vesicle motion with the diffusion model and the inverse relationship between the diffusion coefficient and the strength of the actomyosin network. In active diffusion motion the vesicle diffusion length depends on the mechanical properties and dynamics of the cytoskeleton network in the cells so every object in the cell is expected to move in a similar manner, regardless of its size [37–40, 61]. Therefore, the lack of correlation between the diffusion coefficient and vesicle size (Fig. S4A, B) supports the active diffusion motion modality. Furthermore, the values we measured are similar to the diffusion coefficients measured in mice synapses where the vesicle size is much smaller (~50-100nm) [69]. The lower speed and diffusion coefficient of the vesicles in the ectoderm, compared to the skeletogenic cells, are inversely correlated to the higher signal and activity of the actomyosin network in the ectodermal cells. The strength and activity of the myosin network were shown to define the stiffness of mouse fibroblast [70]. Thus, the higher activity of the actomysin network could constrain the vesicles motion in the cells, and explain the slower speed and lower diffusion length in the ectodermal cells compared to the skeletogenic cells. Furthermore, VEGFR inhibition that completely blocks skeletogenesis, hardly affects the vesicle kinetics and the actomyosin signal. This indicates that vesicle motion is not regulated by VEGF signaling but, mostly likely, is defined by the actomyosin network contractility. Overall, our studies suggest that calcium vesicles undergo active diffusion motion that reflects the mechanical properties of cells and therefore varies between the mesenchymal skeletogenic cells and the epithelial ectodermal cells of the sea urchin embryo.

Our analysis of vesicle movement near the spicules shows that the vesicle motion is not directed towards the spicule but they slow down at distances of 1-2μm from the spicule, as they possibly bind to it (Fig. 6). Furthermore, the spicule is surrounded by actin filaments that coat the membranes around the calcite spicule (Fig. 7, [67]). In the *Drosophila* salivary gland a regulatory network of cytoskeleton proteins, including f-actin, assembles around the vesicles once they bind to the apical membrane and controls the secretion of the vesicle content [63, 64]. This network includes the small GTPase, Rho1, the Rho kinase, ROK, and a RhoGap protein that inactivates Rho1 [64]. Relatedly, inhibition of ROK homolog, inhibition of the small GTPase CDC42 and knockdown of the RhoGap gene, rhogap24l/2 perturb spicule formation [18, 71, 72], supporting an active role of the cytoskeleton machinery in sea urchin skeletogenesis. Possibly, the calcium vesicles attach to the inner membrane of the spicule chord, and secret their content by regulated actomyosin contractions around the vesicle. Further studies of the role of cytoskeleton remodeling proteins in vesicle deposition in the sea urchin embryo will hopefully illuminate the mechanism that control calcium vesicle attachment and secretion into the spicule.

Overall, our findings indicate that calcium bearing vesicles are not performing a directed motion, neither in the cells nor near the spicule. Vesicle motion is largely unaffected by VEGFR inhibition, that is a key regulator of skeletogenesis in the sea urchin embryos. Hence, while the calcium concentration in the vesicles is uniquely regulated in the skeletogenic cells [11, 14, 15], calcium vesicle movement does not seem to be specifically regulated in these cells. Calcium vesicles in the skeleteogenic cells diffuse just like calcium vesicle in non-calcifying cells, with higher diffusion coefficient that correlates with the lower strength and activity of the actomyosin network. As intracellular mineral-bearing vesicles participate in the biomineralization process in various phyla [73–77], the diffusive motion observed here could be relevant to other biomineral-forming organisms.

The absence of directed vesicle motion is quite promising for the design of artificial systems that try to mimic the ability of biomineralizing cells to control mineral properties and growth [5, 78]. In these systems, synthetic vesicles with controlled lipid, protein and mineral content are fabricated. The synthetic vesicles are investigated in search for novel therapeutic approaches for promoting calcification on one hand, and preventing ectopic calcification on the other hand [5, 6, 78]. If mineral bearing vesicles are not actively transported to the biomineralization site, but just perform a diffusive motion, there is one less molecular mechanism to worry about when constructing artificial vesicle-based biomimetic systems. Furthermore, our studies imply that vesicle motion can be altered by changing the mechanical properties of the cells and open a new approach for controlling vesicle dynamics in artificial systems.

## Methods

### Adult animals and embryo cultures

In our studies we used two sea urchin species, the Atlantic species, *Lytechinus variegatus* (*L. variegatus*) and the Mediterranean species *Paracentrotus lividus* (*P. lividus*). Adult *L. variegatus* were obtained from the Duke University Marine Laboratory (Beaufort, NC, USA). Spawning was induced by intracoelomic injection of 0.5M KCl. Embryos were cultured in artificial sea water at 23°C. Adult *P. lividus* were obtained from the Institute of Oceanographic and Limnological Research (IOLR) in Eilat, Israel. They were then kept in aquaria located in a dedicated room, raised in 36ppt in artificial sea water (ASW). Eggs and sperm were obtained by injecting adult sea urchin with 0.5M KCl. Embryos were cultured at 18°C in 0.2μ filtered ASW.

### Calcein staining

A 2mg/ml stock solution of green calcein (C0875, Sigma, Japan) was prepared by dissolving the chemical in distilled water. Working solution of 25μg/ml was prepared by diluting the stock solution in artificial sea water. Embryos were grown in calcein artificial sea water from fertilization and washed from calcein about 2-3 hours prior to the experiments.

### FM4-64 staining

A 100μg/ml stock solution of FM4-64 (T13320, Life technologies, OR, USA) was prepared by dissolving the chemical in distilled water. Working solution of 5μg/ml was prepared by diluting the stock solution in artificial sea water. Calcein stained embryos were immersed in working solution about 10 minutes before visualization.

### Axitinib (AG013736) treatment

A 5mM stock solution of the VEGFR inhibitor, axitinib (AG013736, Selleckchem, Houston, TX, USA), was prepared by reconstituting this chemical in dimethylsulfoxyde (DMSO). Treatments were carried out by diluting aliquots of the axitinib stock in embryo cultures to provide a final concentration of 150nM. Control embryos in all experiments were cultured in equivalent concentrations of DMSO at no more than 0.1% (v/v).

### Sample preparation for Lattice Light Sheet Microscopy

2% low melting agarose (Sigma cat# A0701) melted in artificial sea water at 37°C was added to the stained embryos at the ratio of 5:1, to immobilize the embryos. The sample was then immersed in the microscope tab with 8 mL artificial sea water with 20μL FM4-64 working solution.

### Lattice Light Sheet Microscopy

The lattice light sheet microscope (LLSM) used in these experiments is housed in the Advanced Imaged Center (AIC) at the Howard Hughes Medical Institute Janelia research campus. Prior to image acquisition, the instrument is aligned and operated as previously described [47]. For these samples, a maximum NA of 0.4 and a minimum NA of 0.35 are used to generate the light sheet pattern. It is important to note that due to minute difference in alignment, each LLSM may require different parameters optimization to achieve the highest image quality. Samples are illuminated by a 2D optical lattice (Bessel lattice) generated by a spatial light modulator (SLM, Fourth Dimension Displays). The sample is excited by 488 nm, diode lasers (MPB Communications) at 30mW through an excitation objective (Special Optics, 0.65 NA, 3.74-mm WD). Additional imaging details are included in supplementary dataset S1. Fluorescent emission is collected by detection objective (Nikon, CFI Apo LWD 25XW, 1.1 NA), and detected by a sCMOS camera (Hamamatsu Orca Flash 4.0 v2). Acquired data are deskewed as previously described [47] and deconvolved using an iterative Richardson-Lucy algorithm (10 iterations) with a point-spread function empirically determined for the lattice-light sheet optical system. See the section on data and source code availability for links to sample data and deconvolution source code.

### Segmentation of Calcium Vesicles

The automatic detection of vesicles in each frame (segmentation) is accomplished using the high-performance Hydra Image Processing library for fast 3-D image filtering [79]. We use a blob-detection approach related to Chenourd et al. based on the Laplacian of Gaussians (LOG) filter [51]. The LOG filter can be used for blob-like object detection. In this work we used a thresholded 3D LOG to identify the vesicles in each frame. A scale-invariant anisotropic LOG filter was developed to account for the anisotropic depth dimension of LLSM and other 3D microscope modalities. Specialized edge handling was also developed to allow for consistency in image processing (using LOG and other filters) near the volume edges, this is a key issue when imaging regions of a large organism. The blob-detection method identifies the size and position of each vesicle, some example vesicle detections are shown in figure 4B, based on the image data in Fig. 4A.

### Tracking of Calcium Vesicles

In order to keep track of each vesicle over time, we integrated our segmentation approach into the LEVER software package [52, 53]. LEVER uses a multitemporal cost function based on a small motion model to link the most likely vesicle detections into tracks (Fig. 4B). As part of this work a new bidirectional cost analysis was implemented to improve tracking robustness while allowing faster tracking of vesicles. We used these tracks to analyze vesicle dynamics such as changes in velocity or size over time. The performance of the tracking algorithm degrades with significant motion, beyond the capabilities of the software to predict [52, 57]. In these datasets it is difficult to determine the frame-to-frame vesicle identities even by eye (see *e.g.* movie S8). For this reason, we have restricted our tracking-based analyses to datasets with a time-lapse of 6.12 seconds or less, between consecutive frames.

### Single-Frame Manual Identification of Ectodermal Region

In order to examine the dynamics of calcium specifically in the skeletogenic cells, we must separate out the ectoderm cells which form the exterior layer of the sea urchin embryo. However, in these images, the cells of the ectodermal region are often similar in their general shape to the skeletogenic cells and are quite close together. For this task we manually defined the ectodermal boundary in the first frame of each dataset. An example of such a boundary is shown in figure 4C.

We also applied a registration algorithm using normalized covariance to automatically detect motion of the ectoderm region over time. For all movies analyzed, the motion of the ectoderm region was minimal for the first 50 frames. We therefore limited the analysis of our datasets to the first 50 frames, allowing the use of the ectoderm mask for identifying vesicles throughout the analysis.

### Vesicle Speed, Directionality, and Motion

We measure the frame-to-frame speed of each vesicle by measuring the distance moved between frames divided by the time between frames. This instantaneous speed is averaged over each vesicle track to produce an **average instantaneous speed** per vesicle. Short vesicle tracks (less than 7 frames) are ignored as they are likely to be made up of noisy or unreliable detections.

We also measure the **directionality index** for each vesicle (Fig. 5B). This is the ratio of the maximal displacement of the vesicle divided by the total path length traveled by the vesicle in a window of 60 seconds. For example, in figure 5B, the directionality index ratio would be the length of the white dotted segment (maximal displacement) divided by the total length of the magenta segments. The index is related to the directedness of motion observed for each vesicle, a vesicle traveling in a straight line will have a directionality index of 1, whereas an undirected vesicle will have a low directionality index.

In order to identify the types of motions exhibited by the calcium vesicles, we fit a standard diffusion motion model to each track. For the diffusion model, we measure the **diffusion coefficient** as the slope of the linear relationship between vesicle mean-square displacement (MSD) and time (Fig. S3). All motion and size comparisons were based on the vesicle segmentation and tracking information from the LEVER software, they were computed using MATLAB version 2019b analysis software.

### Spicule-Relative Measurements

Understanding the dynamics of calcium vesicles in biomineralization requires identifying the motion of vesicles relative to the skeletal structure (spicule) in DMSO image sequences. For a selection of movies with a visually defined spicule, we manually identify the spicule centerline in a single frame. An example centerline is shown along with the 3-D spicule image in figure 6A. All vesicle distances are measured relative to the spicule centerline in each frame. Vesicles are binned in 1μm increments by their average distance from the spicule for comparison of average speed at different distances from the spicule. Average distance change per second is also measured for each vesicle, this is referred to as the average velocity toward the spicule.

### Statistical Analysis

For the analysis of differences in average vesicle size and motion characteristics, we first apply the Kruskal-Wallis one-way rank test to identify if there are significant differences across all experimental groups and regions. If a significant difference is observed, then post-hoc pairwise analysis (Dunn-Sidak) is carried out between groups to identify differences between each pair of interest. The MATLAB version 2019b software package was used to perform the analysis. The p-values of each comparison are presented in Dataset S1.

### Phalloidin labeling and 6a9 Immunostaining procedure

Phalloidin labeling and 6a9 immunostaining were done similarly to [66] with minor modifications. Embryos were fixed in 4% paraformaldehyde, 33mM Maleic acid pH7 and 166mM NaCl, for 30 minutes at room temperature. Embryos were washed three times with PBST, then incubated in blocking solution (PBST and 4% sheep serum) for one hour at room temperature. The embryos were incubated overnight at 4°C with 6a9 antibody [65] in blocking solution (1:3-1:5 dilution). The embryos were washed three times in PBST, and then incubated with the secondary antibody (488-Goat anti Mouse IgG 1:200 in PBST Blocking) for 40-60 minute in room temperature. This was followed by 3 PSBT washes, 2 PBS washes, incubation with 50mM ammonium chloride in PBS for 5 min, 3 washes in PBS, incubating with blocking solution (PBS with 4% serum) for 10 minutes in room temperature. Embryos were then treated with hyaluronidase [750 U/ml in PBS (Sigma H3506)] for 10 min, followed by two washes in PBS and one hour in PBS-blocking solution. Embryos were incubated overnight with 1:20 phalloidin-584 (A12381, ThermoFisher) at 4°C, then washed three times in PBS, incubated in PBS-blocking solution for 20 minutes and washed again in PBS.

### MyoIIP and 6a9 Immunostaining procedure

Immunostaining of MyoIIP (p20-MyosinII) and 6a9 were done similarly to [80] with minor modifications. Embryos were fixed in 4% paraformaldehyde as described above and then washed four times with PBST. Embryos were incubated for one hour in PBST-blocking solution, followed by incubation with primary antibody against MyoIIP (Anti-Myosin light chain (phospho S20) Abcam -ab2480. 1:200 dilution in PBST-blocking solution), and 6a9 antibody (1:3 to 1:5 dilution) overnight, at 4°C. Embryos were then washed four time in PBST, then the secondary Antibodies (488-Goat anti Mouse IgG, 115-545-166 and Cy3-Goat Anti Rabbit IgG, 111-165-144 Jackson ImmunoResearch, respectively) were added to the embryos, diluted 1:200 in PBST-blocking solution and incubated for 40-60 minutes in room temperature. This was followed by four washes with PBST and transfer to store solution (PBST in 50% glycerol) at 4°C.

### Phallodin and MyoIIP imaging and quantification

Embryos were imaged in Nikon A1-R Confocal microscope. Three biological replicates (different sets of parents) were imaged for each label and experimental condition. Quantification of phalloidin signal was done for n=7, 9 and 8 control embryos and n= 12, 8 and 10 VEGFR inhibited embryos. Quantification of MyoIIP signal was done for n=8, 7 and 12 control embryos and n= 11, 10 and 9 VEGFR inhibited embryos.

A simple graphical program was written to allow manual identification of ectodermal and skeletogenic cell regions in the phalloidin and MyoIIP images. For each image, at least one ectodermal and skeletogenic region was selected, corresponding to one or more complete cells (Figs. 7G, 8G). A global threshold was chosen for each image to identify the “stained” phalloidin or MyoIIP pixels. For each region the total area was computed along with the area corresponding to the “stained” pixels. The ratio of stained phallodin or MyoIIP to region total area was compared between ectodermal and skeletogenic cells, and between control and VEGFR inhibited embryos. The Kruskal-Wallis one-way rank test was used to compare for overall differences across all groups, followed by pairwise post-hoc (Dunn-Sidak) comparison to identify differences between the pairs of interest. The graphical program and statistical analysis were carried out using MATLAB R2019 software.

### Dataset and Source Code Availability

Sample datasets have been made available at: https://doi.org/10.5281/zenodo.4382712. Two deconvolved lattice light-sheet datasets (100 frames each) from the live-cell experiments are available, one from a control embryo and one from a VEGFR inhibited embryo. Both datasets were used for size and motion statistics in this work. Additionally, the control embryo dataset has also been uploaded in raw form (without deskew or deconvolution). Finally, four confocal datasets from the cytoskeletal remodeling experiments are included, two phalloidin stained images (control/VEGFR inhibited) and two MyoIIP stained images (control/VEGFR inhibited).

Source code for all segmentation, tracking and statistical analyses of the lattice light-sheet data and the graphical tool used for analysis of the phalloidin and MyoIIP datasets are available in the Gitlab repository at: https://git-bioimage.coe.drexel.edu/opensource/llsm-calcium-vesicles-lever. The lattice light-sheet analysis software also requires the LEVER software tools to be installed, available at: https://git-bioimage.coe.drexel.edu/opensource/leverjs. Instructions for installation and example usage of the tools are also provided in the repository readme. Code for the deskew and deconvolution algorithms are available from Janelia at: https://www.janelia.org/open-science/lattice-light-deconvolution-software-cudadeconv.

## Acknowledgements

We thank the advanced imaging center at Janelia research campus for their instrumental help with using the LLSM and in animal maintenance. Specifically, we thank Teng-Leong Chew and John Heddleston for the help with the LLSM, Satya Kuhn for help with animals and reagents and Eric Wait for helpful discussions regarding data analysis. We thank Charles Ettensohn for the generous gift of the 6a9 antibody produced in his lab. We thank Dedi Meiri for the gift of phalloidin. We thank Yarden Ben-Tabou de-Leon for the illustration of the graphical abstract. This work was supported by the Israel Science Foundation grant numbers 41/14 and 211/20 (S.B.D.), Zuckerman fellowship (M.R.W.) and ISEF fellowship (M.M.).

## Author contributions

S.B.D., M.M. and T.G. designed the project. M.M. and S.B.D. performed the LLSM experiments. A.C. and M.R.W. developed visualization, tracking and segmentation tools. M.R.W. analyzed the LLSM data. S.B.D., M.R.W., M.M. and T.G. interpreted the analyses results. T.G. performed the immunostaining experiments. M.R.W. developed a quantification tool that he and T.G. used to quantify the phalloidin and myoIIP signal. S.B.D. and M.R.W. wrote the paper with significant help from M.M. and T.G. and A.C.

## Author information

The authors declare no competing financial interests.

## Notes

https://doi.org/10.5281/zenodo.4382712

https://git-bioimage.coe.drexel.edu/opensource/llsm-calcium-vesicles-lever

### Competing Interest Statement

The authors have declared no competing interest.

### Summary of Updates

This revision has been updated with new experiments and improvements following comments from peer reviewers.

