## Supplemental Text and Figures for "Calcium-vesicles perform active diffusion in the sea urchin embryo during larval biomineralization"

**This PDF file includes:**

Figures and captions S1 to S4

Caption to Dateset S1

Captions to Movies S1-S9

**Other supplementary materials for this manuscript include the following:**

Movies S1-S9

Dataset S1

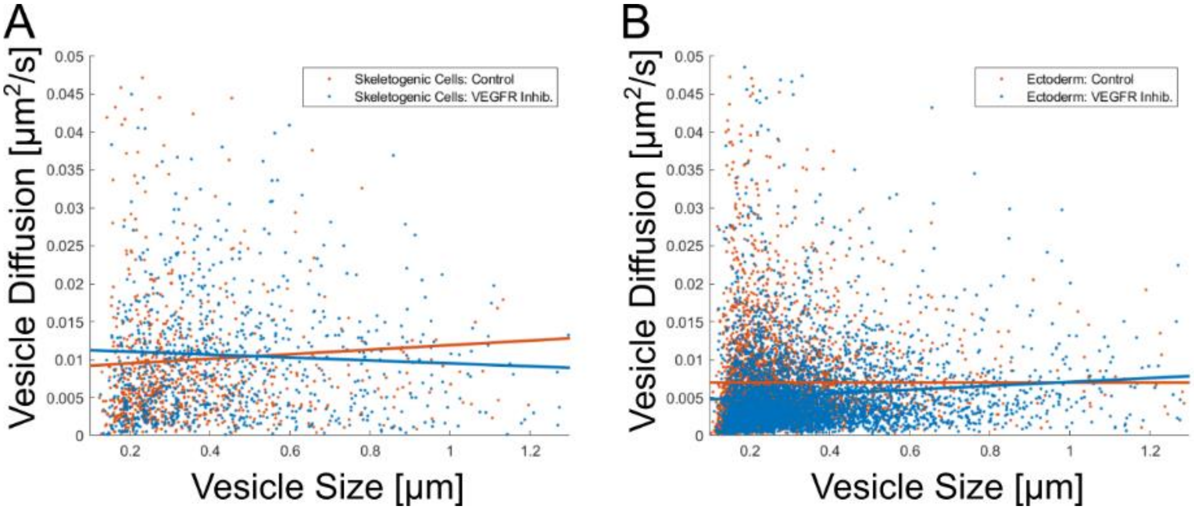


**Supplementary Figure 1 Vesicle instantaneous speed vs. size scatter plot comparison.** These images show average vesicle speed and size for each vesicle tracked, in both skeletogenic mesodermal (A) and ectodermal (B) regions. A trendline fit is shown for each region, in orange for control and in blue for VEGFT inhibition. In both regions and in all experimental condition Pearson's $R^2 < 0.013$, showing no correlation between vesicle size and velocity.

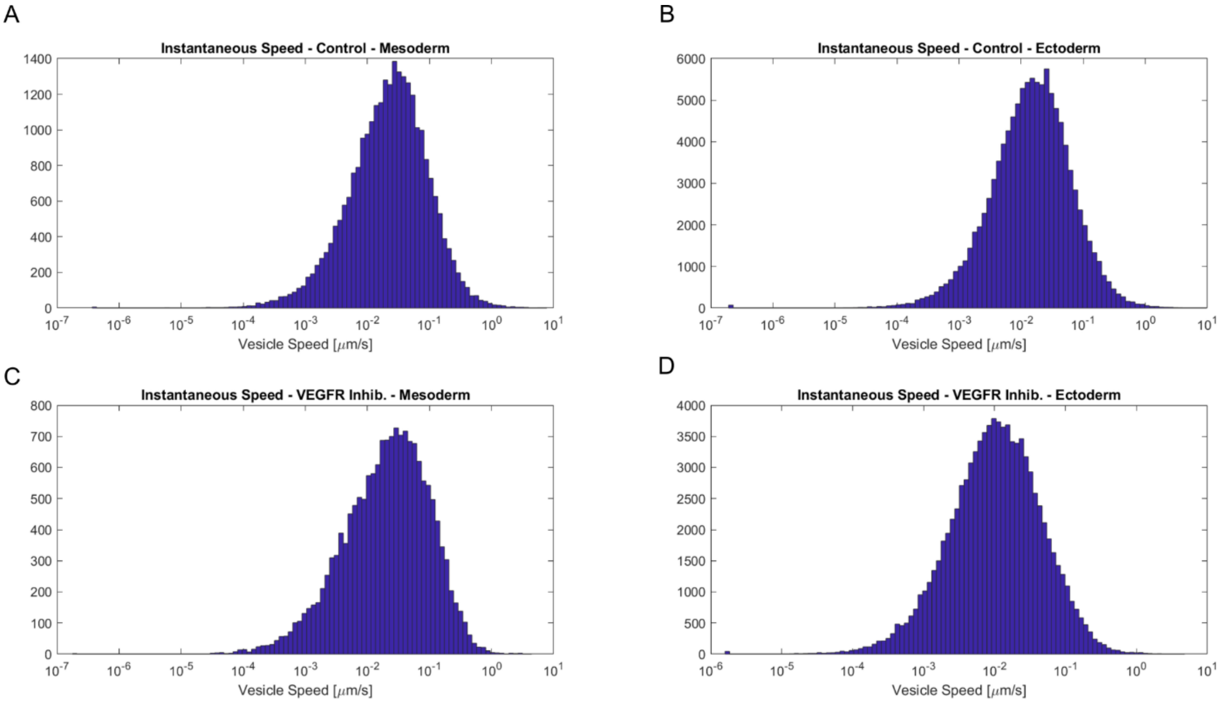


**Supplementary Figure 2 Vesicle instantaneous speed histograms.** Histograms of vesicle instantaneous speed for every track in control (A, B) and VEGFR inhibition (C, D) are shown. For each histogram there is a single peak, making it unlikely that there are multiple different motion types (*e.g.* progressive movement vs. vesicle pausing observed in motor-guided movement).

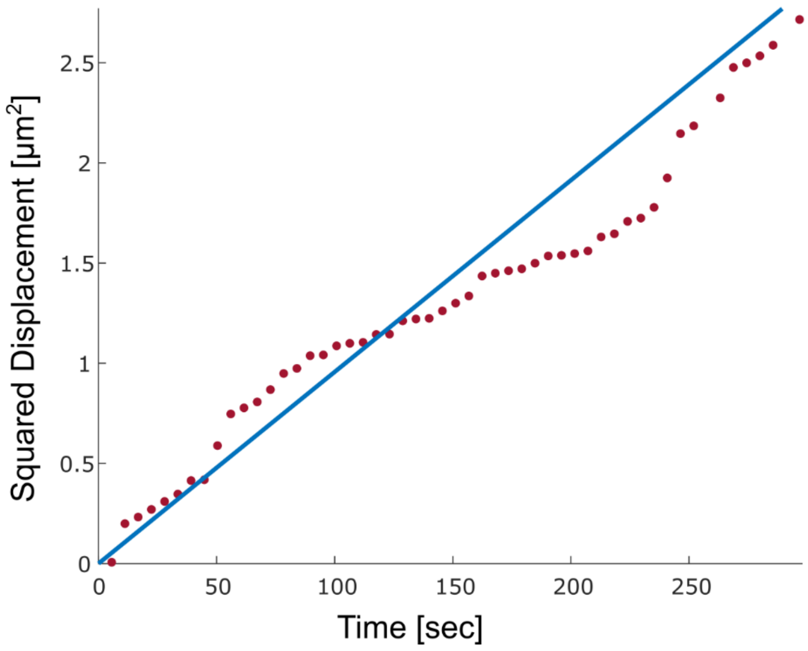


**Supplementary Figure 3 The diffusion coefficient is calculated as the linear fit of the ratio between squared displacement and time.** The plot is an example of this computing the diffusion coefficient as the slope of the linear relationship between squared displacement and time for a single vesicle track.

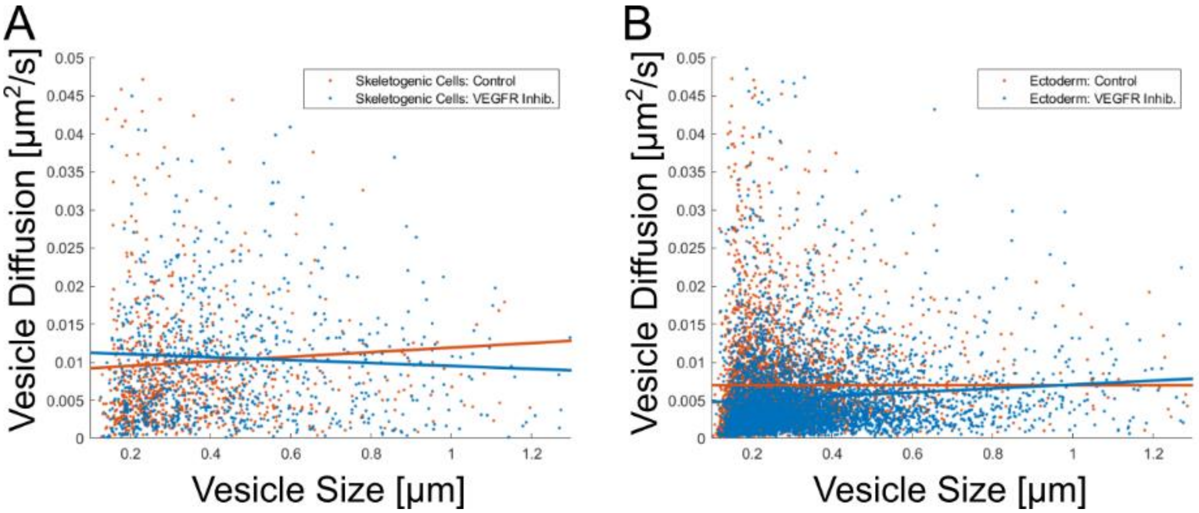


**Supplementary Figure 4 Vesicle diffusion coefficient vs. size scatter plot comparison.** These images show vesicle diffusion rate relative to average vesicle size for each vesicle tracked, in both skeletogenic mesodermal (A) and ectodermal (B) regions. A linear regression fit is shown for each region, orange for control and blue for VEGFR inhibition. In both regions the Pearson's $R^2 < 0.007$, indicating that there is practically no correlation between vesicle size and diffusion rate.

**Dataset S1. Movie analysis reference table and Statistical analysis size and motion results (separate file)**

Spreadsheet 1: This table shows the names of all LLSM datasets analyzed in this work, as well as a note for each dataset that is included as a supplementary movie. Also noted in the table is the temporal resolution of the datasets, whether they were included in motion analysis ($\Delta t$~6.12 sec or less), and the number of ectoderm and skeletogenic mesoderm vesicles used in the size analysis (first-frame) and the motion analysis (total). Spreadsheets 2-7: These spreadsheets contain the statistical comparisons for each size and motion features measured from the vesicle tracks. Comparisons are run across experimental conditions (Control/VEGFR Inhibition) and embryo regions (Ectoderm/Skeletogenic Mesoderm). Each sheet in the dataset contains the comparison of a single measured feature (*e.g.,* size, velocity, etc.) across conditions and regions. In the case of spicule-relative analysis, comparisons are run across groups of vesicles at different average distance from the spicule binned in 1μm groups. The Kruskal-Wallis test is used to establish differences in centrality between groups, followed by a post-hoc Dunn-Sidak test to establish pairwise differences. A brief description of the comparison is included at the bottom of each sheet.

**Movie S1 (separate file)**

Movie showing the a single frame of the 3D lattice light-sheet imaged region of a representative embryo showing both the cell membranes (FM4-64,white) and calcium vesicles (calcein,green). The volume is visualized by perspective projection onto a viewing plane by raycast sampling of the volumetric data. While the data in supplementary movies S2-S7 and S9 are also 3D volumetric raycast projections, we visualize them from a fixed perspective over time, in order to maintain consistency for vesicle and cell motion from frame to frame.

**Movie S2. (separate file)**

Movie showing a control embryo before spicule formation. There is significant motion and interaction among the skeletogenic cells forming a cluster at the bottom of the frame. Calcein staining is shown in green identifying calcium vesicles and FM4-64 membrane staining is shown in white.

**Movie S3. (separate file)**

Movie showing a control embryo with a small spicule. The spicule is clearly visible, surrounded by the skeletogenic cells. Calcein staining is shown in green identifying calcium and FM4-64 membrane staining is shown in white.

**Movie S4. (separate file)**

Movie showing a control embryo with an elongated spicule. The base of the spicule is stained with Calcein and a portion of the length of the spicule can only be discerned by the enclosing tube structure stained with FM4-64 membrane marker. Calcein staining is shown in green identifying calcium and FM4-64 membrane staining is shown in white.

**Movie S5. (separate file)**

Movie showing a VEGFR inhibited embryo. A large cluster of skeletogenic cells is shown in the bottom left of the frame. Calcein staining is shown in green identifying calcium vesicles and FM4-64 membrane staining is shown in white.

**Movie S6. (separate file)**

Movie showing a VEGFR inhibited embryo. A cluster of skeletogenic cells is shown in the bottom left of the frame. Calcein staining is shown in green identifying calcium vesicles and FM4-64 membrane staining is shown in white.

**Movie S7. (separate file)**

Movie showing a VEGFR inhibited embryo. A large cluster of skeletogenic cells is shown in the bottom left of the frame. Calcein staining is shown in green identifying calcium vesicles and FM4-64 staining is shown in white.

**Movie S8 (separate file)**

Movie showing a single frame of example vesicle segmentations overlaid on the volumetric data. The segmentation data includes a triangle mesh (visualized here) representing the closed exterior boundary of each vesicle, as well as computed internal volume and other characteristics, such as center of mass which is used to compute motion statistics.

**Movie S9. (separate file)**

The movie shows the identification and tracking of vesicles over time. The movie focuses on four vesicles highlighted in color overlaid with identity labels. There is significant motion throughout the video, but the vesicles do not appear to be directed in their motion. Colored tails are drawn along the previous centroids of vesicles to show the vesicle’s motion over time.
